# Chemokine Signaling Caused by *Mycobacterium avium* Biofilms in the Lung Airway Increases Bacterial Loads by Spatially Diverting Macrophages

**DOI:** 10.1101/2022.09.13.507811

**Authors:** Catherine Weathered, Kelly Pennington, Patricio Escalante, Elsje Pienaar

## Abstract

*Mycobacterium avium* Complex (MAC) are ubiquitous environmental biofilm-forming microbes that can colonize and infect patient lungs. Incidence and prevalence of MAC infections are increasing globally, and reinfection is common. Thus, MAC infections present a significant public health challenge. MAC infections are notoriously difficult to treat and there is an urgent need for MAC-targeted therapeutics. To identify potential drug targets, we quantify the impact of MAC biofilms and repeated exposure on infection progression using a computational model of MAC infection in lung airways.

MAC biofilms aid epithelial cell invasion, cause premature macrophage apoptosis, and limit antibiotic efficacy. We develop an agent-based model that incorporates the interactions between bacteria, biofilm and immune cells. We perform virtual knockouts to quantify the effects of the sources of biofilm (biofilm simultaneously deposited with bacteria vs. formed in the airway after initial bacterial deposition), and their effects on macrophages (inducing apoptosis and slowing phagocytosis). We also quantify the effects of repeated bacterial exposure to assess the impact of reinfection on infection progression.

Our results show that chemokines released by biofilm-induced apoptosis bias macrophage chemotaxis towards pockets of infected and apoptosed macrophages. This bias results in fewer macrophages finding extracellular bacteria, allowing the extracellular planktonic bacteria to replicate freely. These spatial macrophage trends are further exacerbated with repeated deposition of bacteria.

Our model indicates that interventions to either abrogate macrophages’ apoptotic responses to bacterial biofilms and/or reduce frequency of patient exposure to bacteria will lower bacterial load, and likely overall risk of infection.

## 1. Introduction

*Mycobacterium avium* complex (MAC) is an opportunistic infection of the lungs, and the most common causative agent of non-tuberculous mycobacterial pulmonary disease (NTM-PD). These infections disproportionately affect patients with preexisting lung damage, Cystic Fibrosis, Chronic Obstructive Pulmonary Disease (COPD), and post-menopausal women, and incidence and prevalence are rising^1^. NTM-PD significantly lowers quality of life of patients, and requires prolonged multi-drug antibiotic regimens that last at least 1 year after sputum conversion and are often difficult to tolerate by patients ^2^, with a success rate of only 45-65%^3^. Additionally, patients who have reached a clinical cure for these infections remain at increased risk of reinfection^4^.

MAC infections occur when bacteria are aerosolized, inhaled, and deposited in lung airways. Many potential sources of MAC have been identified, ranging from water sources^5^ to soil^6^. In one study, pathogenic mycobacteria were isolated from water sources within 19 of the homes of 20 NTM-PD patients, and bacterial strains matched in seven of the patients^7^. There is, however, a clinical distinction between airway colonization, where bacteria are present in sputum, vs. infection, which is characterized by clinical and radiographic evidence of airways involvement and nodules in the lungs^8^. These early events and dynamics in the airways are difficult to study experimentally but may play an important role in infection progression. Further, bacterial dynamics in the airways are important to understand, since sputum samples expectorated from the airways are a clinically important measure of infection and treatment efficacy.

One hypothesis to explain the difficulty in treatment is the potential for MAC to exist in biofilms *in vivo* in the lungs, as they are known to do in the environment. *In vitro*, MAC biofilms have been shown to decrease bacterial susceptibility to antibiotics^9^, increase bacterial invasion into epithelial cells^10^, prevent or decrease macrophage phagocytosis of bacteria^11^, and cause premature apoptosis of macrophages^11^. In this work, we decouple the sources and effects of biofilms to quantify how they aid bacteria or hinder immune cells; and we determine the impact of single exposure vs. repeated bacterial exposure on infection progression. Thus, we can better identify potential interventions or prophylactic approaches to prevent recurrent infection.

We take a systems biology approach, using an established agent-based model of host macrophage-bacterial interactions in the lung airway^12^. This spatio-temporal model allows us to examine interactions between bacteria, their biofilms and macrophages over the initial two weeks post-inhalation. First, we investigate the role of biofilm deposited with the initial bacterial inoculum compared to biofilm contributed by bacteria within the lung airway after inhalation. We probe these differences by performing knock-out experiments on both potential sources of biofilm separately. This serves as a marker for the best-case scenario in the use of an antibiofilm intervention. Second, we examine the effects of modulating host immune responses to biofilm, which will inform host-directed therapy targets to ameliorate the impact of biofilm on the host immune response. Third, we examine the impact of repeated inhalation events by introducing new bacteria to the lung airway at regular intervals while the host macrophage-bacterial interactions are already underway. We evaluate the distribution of bacterial phenotypes as new planktonic and sessile bacteria are successively added to the airway environment.

## 2. Methods

### 2.1 Model Overview

Our agent-based model, developed in Repast Simphony^13^, describes the interactions between MAC bacteria, their biofilms and host macrophages in the lung airway^12^ (Figure 1a). Briefly, a mixture of planktonic and sessile bacteria is deposited in the airway. This deposition triggers macrophage chemotaxis and phagocytosis of bacteria. Biofilms can be both deposited with sessile bacteria or contributed by sessile bacteria after deposition in the airway. Once formed, biofilms can cause macrophages to become overstimulated and undergo apoptosis^11^.

**Figure 1.**
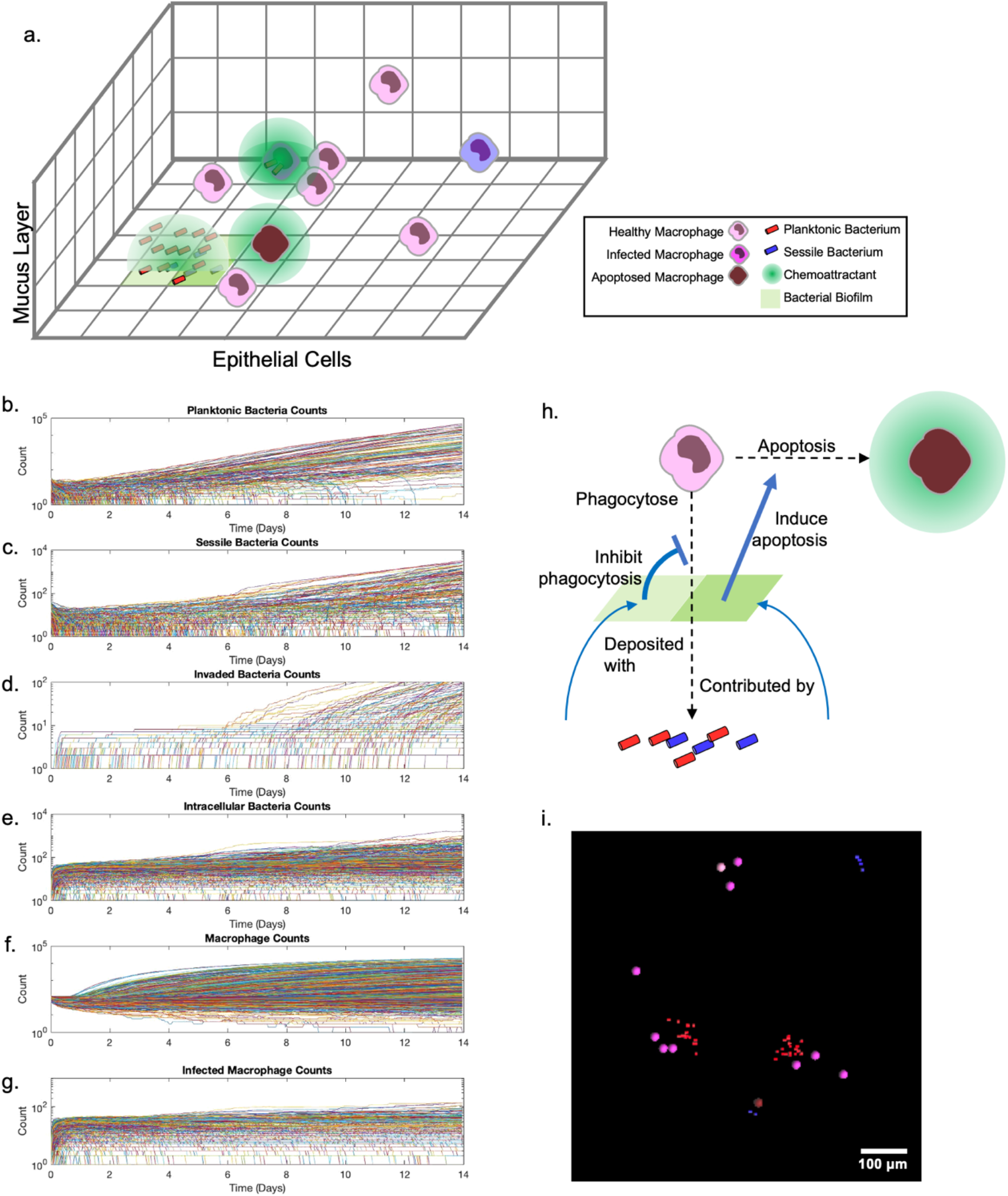
Model Diagrams, timeseries outputs, and representative simulation. a. Agent interactions take place in a three-dimensional grid which represents the lung airway, with toroidal boundaries in the x and y dimensions and epithelial cells at the bottom of the z-dimension. b-g. Timeseries counts of each cell subtype are recorded for each simulation. Here we show the control timeseries counts for b. planktonic, c. sessile, d. invaded, and e. intracellular bacteria, as well as (healthy) macrophages and infected macrophages in each of our 900 control simulations. h. Our mechanisms of interest in this paper are the interactions of bacteria with biofilms and biofilms with macrophages. Bacteria contribute biofilms to the environment by either being deposited with pre-formed biofilms or producing new biofilm once in the lung airway. These biofilms slow or prevent macrophage phagocytosis of bacteria within the biofilms and cause apoptosis. i. When these interactions are allowed to unfold in the simulation environment, spatially heterogeneous infections emerge. Image shows 750 by 750 um portion of a simulation space 7 days after exposure.

The model is executed in six-minute timesteps for a total of 14 simulation days. Model outputs include timeseries counts of each cell subtype (**Error! Reference source not found**.b-g). Timeseries data are collected as spatially heterogeneous infections emerge (Figure 1i) that can, in turn, be probed for other data such as spatial chemokine concentrations. A complete description of the model can be found in Weathered et al.^12^, but mechanisms relevant to this work (Figure 1h) are briefly described below.

### 2.2 Macrophage recruitment

Macrophages are recruited to the lung airway via grid compartments designated as “recruitment areas” spread randomly throughout the airway. In previous model iterations, once a chemokine threshold is reached at a given recruitment site, a constant probability (sampled within a range) is used to determine if a macrophage would be recruited at that timestep. The implementation therefore resembled a step function. This mechanism is updated here to include a more biologically relevant continuous function^14,15^. The model now uses a likelihood function of recruitment that increases with chemokine concentration, but recruitment is possible at any concentration. This is implemented as a Hill function (Supplement 2), in which β is the maximum probability of recruitment, K is the chemokine concentration where half of the maximum recruitment probability is reached, and n represents the sensitivity of recruitment to chemokine concentration. A full parameter table is in Supplement 1.

### 2.3 Biofilms

In the simulation environment, biofilm can be introduced in two different ways. Biofilm can be: a) deposited with the initial bacterial inoculum, representing biofilm that sheared off the surface where the bacteria originate, or b) contributed by sessile bacteria after they are established in the environment. We represent biofilm as a continuous spatial variable, where each grid compartment has some amount of biofilm ranging from zero to 100. The amount of biofilm is used to determine the biofilm’s effects on macrophages (discussed below).

When bacteria are deposited in the airway, each sessile bacterium is deposited with *A_itlBiofilm* biofilm in the same grid compartment. Additionally, each sessile bacterium can also contribute biofilm to its grid compartment, at a rate *A_bacToBiofilm*.

### 2.4 Macrophage response to biofilms

Biofilms affect macrophages by inducing apoptosis and slowing or preventing phagocytosis of bacteria residing within biofilms.

At initiation of the model, each macrophage is assigned an individual biofilm tolerance, which was calibrated in Weathered et al. to match the rates of apoptosis of macrophages when exposed to biofilms *in vitro*. In each timestep where a macrophage is exposed to biofilm it loses some of its tolerance, at a linear dose-dependent rate. When tolerance reaches zero, the macrophage undergoes apoptosis. After undergoing apoptosis, dead macrophages remain in the environment for a length of time (*T_apopDeg*), releasing chemokines at the same rate as infected macrophages.

Similarly, macrophage phagocytosis of biofilm-associated bacteria is modeled in a dose-dependent manner. Each timestep, a macrophage will check its Moore neighborhood for bacteria to phagocytose, and preferentially phagocytose a planktonic bacterium not in a biofilm. If there are only sessile bacteria, it will select one, then have a probability inversely proportional to the amount of biofilm (out of 100) in that grid compartment of phagocytosing it. This effectively lowers the rate of phagocytosis of sessile bacteria while not completely preventing their phagocytosis.

### 2.5 Macrophage chemotaxis

Macrophages undergo chemotaxis in response to the total chemoattractant in their Moore neighborhoods, including their current position. This total attractant layer is the sum of the diffusible chemoattractants released by bacteria and macrophages. The concentration of chemoattractant in each grid compartment of the Moore neighborhood is used to probabilistically weight the macrophage in the direction of the higher concentration. If the total concentration is under *A_chemotaxThresh* the macrophage chooses a new grid compartment randomly.

### 2.6 Virtual Knockouts

To assess the impact of biofilm sources, we compare four groups of simulations: 1) control simulations that include both deposited and contributed biofilms, 2) contributed-biofilm knockout simulations, 3) deposited-biofilm knockout simulations, and 4) simulations where both deposited and contributed biofilm is knocked out. Each group consists of 900 simulations: 300 parameter combinations randomly selected from the parameter ranges in Supplement 1, each run in triplicate to account for stochasticity in the model.

We use the same experimental design to knock out macrophage-based mechanisms, eliminating 1) the biofilm’s ability to prevent macrophage phagocytose bacteria in biofilms and 2) macrophage apoptosis in response to biofilms (while maintaining their ability to undergo apoptosis in response to internal bacteria).

Finally, in our chemotactic knock-out simulations, we alter all macrophage movement to be a random walk, as it is when the chemokine concentration is below the macrophage limit of detection.

For statistical analysis, we use a one-way analysis of variance (ANOVA, α=0.01) to compare differences among groups. If the null is rejected, indicating that there is a difference in the mean value of the measure between groups, we follow with a Tukey-Kramer post-hoc test. In testing for statistical equivalence, we use Two One-Sided Test^16,17^ (TOST).

### 2.7 Repeat Deposition

MAC can be deposited into the airway via inhalation of aerosolized bacteria, from local sources including waterways and showerheads. The high reinfection rate of NTM^4^ indicates that re-exposure is common, and the ubiquity of the environmental sources indicates that inhalation is likely not a one-time event. Instead, it is likely that repeated doses of bacteria contribute to a population already colonizing the airway.

To quantify the impact of repeated exposure, we use centered Latin Hypercube Sampling to select 100 parameter combinations from the parameter ranges in Supplement 1, with three replicates each. For each of these 300 simulations, we vary the deposition frequency, keeping the bacterial load for each deposition the same. We evaluate deposition frequencies of every one, two, three-and-a-half, or seven days, and compare infection progression to control simulations of single-deposition.

## 3. Results

### 3.1 Biofilms are not necessary to form an infection but do worsen bacterial load and macrophage infiltration through increased chemokine signaling and excess macrophage recruitment

In knocking out biofilm that is contributed by inhaled bacteria, biofilm deposited in the airway upon inhalation, or both (the “double knockout”), only the deposited biofilm or double knockouts have a significant impact. Our results show a significant reduction in total bacterial load, bacterial invasion, macrophage recruitment, and macrophage apoptosis (Figure 2a,b,d,e) for the deposition knockout and the double knock-out at the end of the 14-day simulation. The deposition and double knock-outs are statistically equivalent to each other (TOST, p=0.493). The importance of deposited biofilm is consistent with previous work^12^, where we showed that the biofilm contributed within the first two weeks post-inhalation by MAC is too slow to have a significant impact on total biofilm. However, the mechanism(s) by which deposited biofilm exacerbate infection progression is not clear from these data.

**Figure 2.**
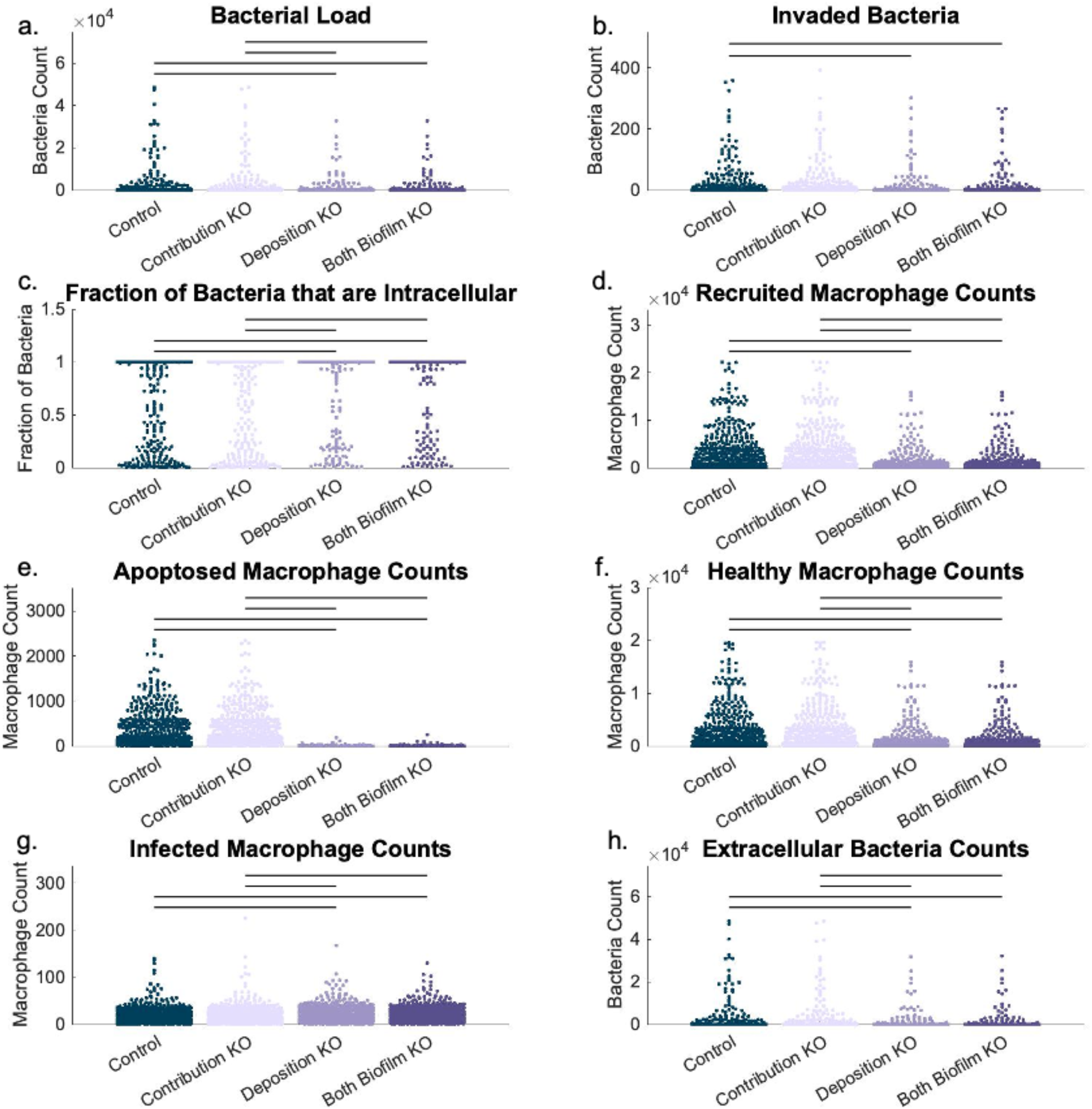
Comparison of control simulations (no knock-outs) to knock-outs of biofilm contributed by bacteria after deposition in the lung airway, biofilms deposited with bacteria in the lung airway, and both biofilm sources. Significance bars indicate p<0.01 in Tukey-Kramer post-hoc test. Each metric is at the end of the simulation, either 14 days or when all bacteria have been cleared. a) Bacterial load is a sum of all planktonic, sessile, and intracellular bacteria. b) Invaded bacteria are those that have moved into epithelial cells, and are no longer counted in the bacterial load. c) Fraction of bacteria that are intracellular is the ratio of intracellular vs. bacterial load, and indicates macrophages’ control of extracellular bacteira through phagocytosis. d) Recruited macrophages are those brought into the lung airway through recruitment areas via macrophage signaling, detailed in section 2.2. e) Apoptosed macrophage counts are those macrophages that have undergone apoptosis, either via biofilm exposure mechanisms or from high intracellular bacteria counts. f) Healthy macrophage counts are those macrophages in the environvment that do not have intracellular bacteria and have not undergone apoptosis. g) Infected macrophages contain intracellular bacteria. h) Extracellular bacteria counts is the sum of planktonic and sessile bacteria in the lung airway.

There is a significant decrease in macrophage recruitment for the deposition knockout, which may be considered beneficial since it reduces infiltrates into the airway. Because there is a corresponding decrease in bacterial load, we can assume that these recruited macrophages are in excess and may actually contribute to inflammation. This reduction in recruitment in the deposition knockout is due to a corresponding decrease in apoptosis (Figure 2e), leading to lower chemokine levels which recruit macrophages. There is a significant increase in infected macrophage counts (those with persistent intracellular bacteria) (Figure 2g) in the deposition knockout. This increase in infected macrophages indicates that the decreased recruitment is not reducing macrophages’ ability to effectively contain the extracellular bacterial population (Figure 2h). Instead, the reduced recruitment only lowers the number of healthy macrophages that are not directly involved in phagocytosing and killing bacteria but could contribute to overall inflammation and infiltrates in the airway.

Taken together, we show that while deposited biofilms do increase bacterial loads and macrophage infiltration, they are not necessary to establish an infection. The simulations with all biofilm sources knocked out also had high numbers of simulations where bacteria were not cleared and with invasion into epithelial cells. Overall, we observe that knocking out biofilm that is deposited with sessile bacteria leads to lower bacterial loads and decreased invasion into the epithelium. We also see decreased apoptosis and decreased recruitment of superfluous macrophages due to increased chemokine release. These data show that targeting biofilm can reduce the number of macrophages being recruited, without compromising the ability of macrophages to contain the bacteria. However, the specific mechanism by which the biofilm knockout results in decreased bacteria remains unclear. Knowing the importance of the effects of biofilms on macrophages, we next examine the ability of biofilm to prevent macrophage phagocytosis of bacteria (protective to bacteria) or cause macrophage apoptosis (offensive against macrophages).

### 3.2 Despite recruitment maintaining macrophage counts, apoptosis results in an increase in bacterial load

To simulate pharmacologically targeting macrophage pathways rather than bacterial biofilm pathways, we focus on two mechanisms via which biofilms affect macrophages: inducing apoptosis and preventing phagocytosis of bacteria within biofilms. Our simulations show a significant reduction in bacterial load (Figure 3a, p=0.006) when the macrophages’ apoptotic response to biofilm is knocked out. Notably, the reduction in bacterial load is statistically equivalent between the macrophage apoptosis knock-out and the deposited biofilm knock-out (TOST, p=0.571), indicating that either intervention may be similarly effective.

**Figure 3.**
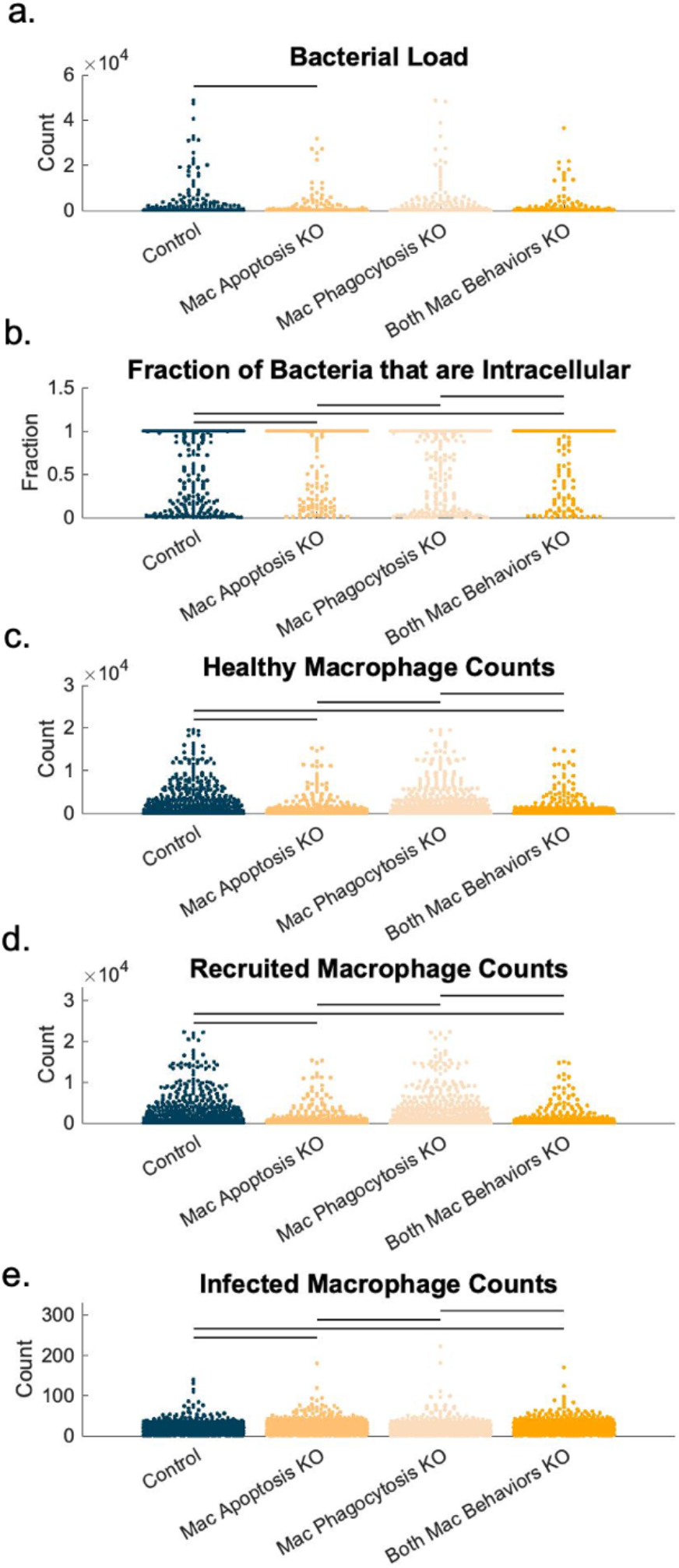
Comparison between control simulations (no knock-outs) to mechanistic knock-outs of macrophage apoptosis in response to biofilms, macrophage phagocytosis being prevented by biofilms, and both. Significance bars indicate p<0.01 in Tukey-Kramer post-hoc test. Metrics are the same as described in Figure 2 caption a, c, f, and g.

Across all metrics, there is no significant effect for knocking out the phagocytosis prevention mechanism. With more mature, fully formed biofilms this mechanism may play a larger role, but in our simulations phagocytosis rates themselves are rarely the limiting factor in macrophages’ response to bacteria.

There is also reduction in mean healthy macrophage counts (Figure 3c) and recruitment (Figure 3d), and a corresponding increase in infected macrophage counts (Figure 3e) in the macrophage apoptosis or combined knock-out simulations. Though the decrease in healthy macrophages may indicate a problem, the significant increase in fraction of bacteria that are intracellular (Figure 3b, p<0.0001) indicates that the macrophages overall control the extracellular bacteria more efficiently with the reduced apoptosis.

Knowing that macrophage apoptosis prevents macrophages from controlling extracellular bacteria efficiently, and not just because it reduces macrophage numbers through killing, we next examine the spatial role of apoptotic chemokine signaling.

### 3.3 Spatial heterogeneity demonstrates the importance of macrophage chemotaxis in finding extracellular bacteria

Bacteria remain in the extracellular space, despite large numbers of healthy macrophages in the airway. Further, prevention of phagocytosis by biofilm is not playing a significant role in either bacterial load or fraction of bacteria that are intracellular. These findings suggest that it is neither macrophage numbers nor rate of phagocytosis that is the limiting factor in controlling extracellular bacteria. We hypothesize that the limiting factor in macrophages’ ability to internalize bacteria is largely finding them. Macrophages must collocate with the bacteria to phagocytose them. To understand how macrophages find both extracellular bacteria and infected or apoptosed macrophages, we examine the chemotactic gradients that drive macrophage movement.

We have previously shown that most phagocytosis events occur early in the simulation before new healthy macrophages are recruited^12^. Macrophages’ ability to find sites of infection depends on chemotactic responses to chemokine gradients. To quantify how effective chemotaxis is in finding the sites of infection in the airways, we knock out macrophages’ ability to undergo chemotaxis, instead forcing all macrophages to move in a random walk. At 24 hours post-deposition, our results show an increase in bacterial load (Figure 4a) and smaller fraction of intracellular bacteria in the chemotaxis knock-out (p<0.0001) (Figure 4b). This indicates that chemotaxis does play an important role in macrophages collocating to bacteria early in the simulation. However, by the end of the simulation there is no significant difference between the control and the chemotaxis knock-out in either bacterial load or fraction of intracellular bacteria (Figure 4c-d). This indicates that the efficacy of chemotaxis in leading macrophages to extracellular bacteria diminishes over time. We hypothesize that this diminished impact of chemotaxis over time is due to excess chemokine signaling of infected or apoptosed macrophages, which may preferentially lead macrophages towards these macrophages and away from extracellular bacteria. Because much of this excess chemotactic signaling is due to biofilm-induced-macrophage apoptosis, we next examine how eliminating the biofilm’s effects on the host immune cells (rather than eliminating the biofilm itself) might impact colonization and infection.

**Figure 4.**
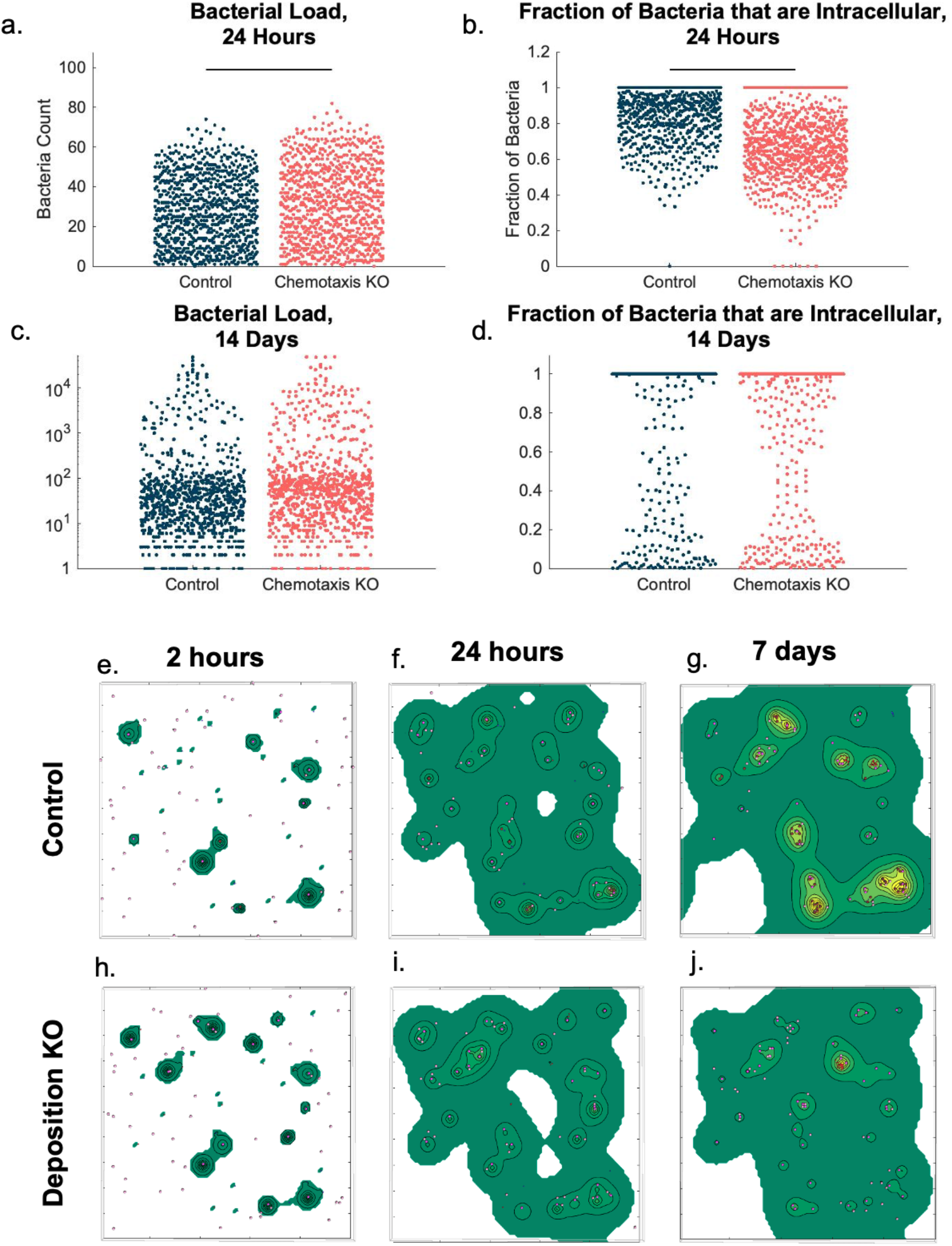
Chemotactic gradient effects. a-b) Beeswarm plots of the total bacterial load and their intracellular fractions at 24 hours between the control and a mechanistic knock-out of macrophage chemotaxis. In the knock-out, macrophages undergo a completely random walk rather than following gradients. c-h) Representative samples between corresponding control and deposited biofilm knockouts at 2 hours, 24 hours and 7 days post initiation. In d-e, we can see significantly more areas of high levels of chemokine gradients, which pull more macrophages to that area. In contrast, h shows that the highest area of chemokine concentration is around an extracellular colony of bacteria, which will eventually guide more macrophages to that area.

### 3.4 Reducing chemokine signaling due to apoptosis aids in collocation of macrophages to bacteria and increased phagocytosis

In both the biofilm deposition knockout and apoptotic response knockout, we saw not only a reduction in bacterial load (Figure 2a and Figure 3a, respectively), but a significant shift on average to a more intracellular phenotype (p<0.0001 for both in comparison to control) (Figure 2c and Figure 3b). After the initial bout of macrophage chemotaxis and phagocytosis following bacterial deposition, many macrophages are either infected or have undergone apoptosis. These infected or apoptosed macrophages are continuously releasing chemokines. Though bacteria are also continuously releasing chemotactic molecules, the host-derived chemokine release creates a much further-reaching chemotactic gradient in our model than that produced by the bacteria.

This gradient causes most macrophages, either those already in the environment or those recruited by the host-derived chemokines, to move towards the infected and apoptosed macrophages, rather than locating and phagocytosing remaining extracellular bacteria. This bias in chemotaxis can also lead to a positive feedback loop, with more recruited macrophages being exposed to biofilm, undergoing apoptosis, and contributing more chemokine signals.

These spatial influences can be seen in representative simulations (Figure 4c-h), where the sum of the chemoattractant layers is represented by a heatmap, overlaid on the spatial map of agents. We can see a wider sphere of chemoattractant around apoptosed macrophages than extracellular bacteria, especially in early timepoints, where their signal is completely lost in the noise of other gradients. It is not until the colony has grown to include several extracellular bacteria that they have their own sphere of chemotactic influence, which may eventually lead more macrophages to them.

In the biofilm deposition knockout, chemokine signaling is reduced because macrophages are less likely to undergo apoptosis, and a higher fraction of the chemoattractant is from extracellular bacteria. We can see this effect in the Deposition KO (Figure 4h), which has fewer apoptotic macrophages (due to the reduction of biofilm) and shows only one high area of chemoattractant, which contains several extracellular bacteria.

These trends show both the influence of spatial factors and excess inflammation. Excess chemokine signaling from either apoptosed macrophages or large groups of infected macrophages can draw all airway macrophages to them, preventing random patrolling and allowing extracellular bacterial colonies to grow unnoticed by macrophages. This problem is compounded when macrophages are recruited because, in our model, the likelihood of recruitment is due to the macrophage chemokine concentration, meaning that the recruited macrophage will most likely already be in a steep chemotactic gradient and most likely to move towards infected or apoptosed macrophages when recruited.

### 3.5 Macrophages being drawn to areas of high chemokine signaling also leads to higher relative bacterial loads for repeated deposition scenarios

So far, we have also been assuming a single deposition of bacteria that then persists over the course of two weeks. Common sources of MAC include soil and showerheads, suggesting that patients are likely exposed repeatedly over the course of our two-week simulations. We hypothesize that the problem of macrophages not finding extracellular bacteria may be further exacerbated when bacteria are added to an already-infected airway. If macrophages are already drawn to a region of higher chemokine signaling and are no longer patrolling the airway, newly, randomly deposited extracellular bacteria may have more time to grow unchecked.

As expected, with repeated deposition of bacteria we observe a significant increase in bacterial load by the end of the simulations (Figure 5a), not only in net bacterial load, but also normalized to the number of bacteria added over the course of the simulation (Figure 5b). Interestingly, there increase in bacterial load is only seen at frequencies of 7x/week (daily) and 3.5x/week (every other day). Every tested frequency lower than that has no significant difference in bacterial load, indicating that there is a threshold at which further reduction does not further decrease bacterial loads.

**Figure 5.**
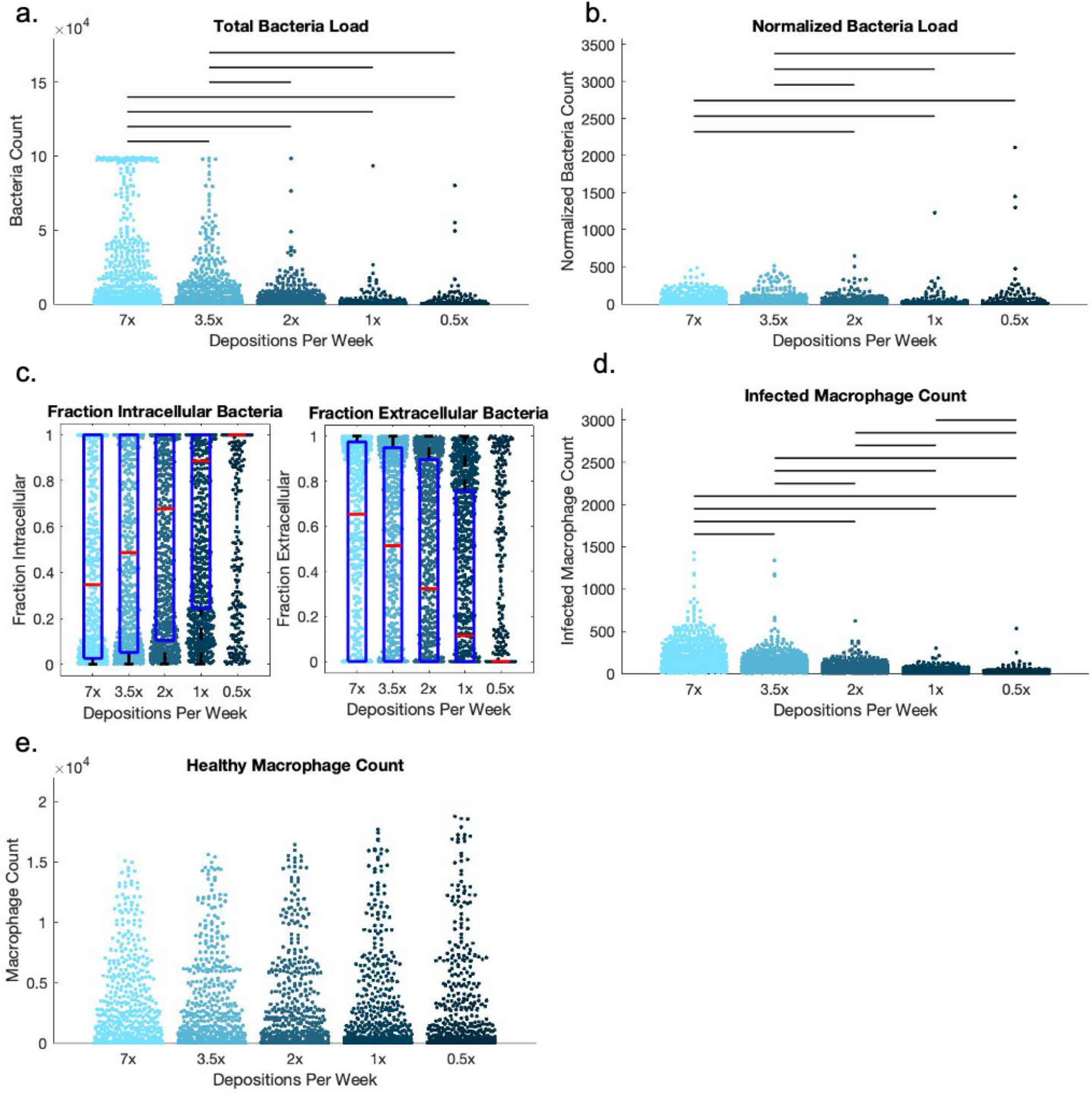
Comparison across frequency of bacterial deposition. In each group, the same initial number of bacteria were deposited either 7x per week (daily), 3.5x per week (every other day), 2x per week (every 3.5 days), 1x week (2x per simulation) or 0.5x per week (our control). Significance bars indicate p<0.01 in Tukey-Kramer post-hoc test.. a) The total bacterial load is understandably higher with more frequent depositions. b) Here the total bacterial load at the end of the simulation is normalized to the total number of bacteria that were deposited over the course of the simulation. We can still see a signfiicant increase in bacteria over the course of the simulation, indicating that increased bacteria deposition is not solely responsible for differences in simulations. c-d) Boxplot overlays on the beeswamrms show corresponding increases in fraction of bacteria that are intracellular and fraction of bacteria that are extracellular as frequency of deposition decreases. e) Infected macrophage counts are signfiicantly higher with more frequent deposition, but f) healthy macrophage counts do not show a significant difference across deposition frequencies.

For each increase in frequency, we see a corresponding increase in infected macrophage counts (Figure 5e), but no significant increase in healthy macrophage counts (Figure 5d). Further, we note a shift to higher proportions of extracellular phenotypes of bacteria (Figure 5c and Supplement 3). This trend is caused by a combination of more extracellular bacteria being added to the airway environment and failure to find and phagocytose these bacteria.

In PRCC analysis, we observe that the initial number of bacteria deposited, which is used for each subsequent deposition, also continues to have a significant impact on total bacterial load (Supplement 4). Because both the frequency of deposition and bacterial load deposited each time are significantly and positively correlated with higher bacterial loads, reducing patient exposure through either frequency or exposure load may reduce overall chance of developing an infection.

## 4. Discussion

In this work we show the role that excess macrophage chemokine signaling plays in allowing a MAC infection to form, and what roles biofilm could play in fueling this inflammation. We perform *in silico* knockouts of both biofilm and macrophage responses to biofilms. With these simulations, we show a significant reduction in bacterial load by knocking out either biofilm that is deposited with sessile bacteria upon inhalation (the primary source of bacterial biofilms in the lung airways in the first 14 days post-exposure), or by knocking out macrophage’s apoptotic response to these biofilms. We demonstrate that the pro-inflammatory response to biofilms cause macrophages to undergo chemotaxis towards other distressed macrophages, rather than extracellular bacteria, and that this problem is exacerbated if more bacteria are regularly deposited in the lung airways.

### 4.1 Preventing or reducing MAC infections through pharmacological interventions

Biofilms are a difficult pharmacological or environmental target. Mycobacterium biofilms specifically have plagued waterways and hospital systems^5,18,19^. However, MAC and other pathogenic mycobacterial biofilm seem to be mainly composed by cellulose in the airways, which could be a potential therapeutic target^20^. However, here we show that eliminating macrophage apoptotic responses to biofilm is just as effective in aiding host response to bacteria as eliminating the biofilms themselves. We believe that in the future, this may be accomplished either pharmacologically, as a prophylactic intervention for patients most at risk (e.g. those with prior infections), or via vaccines, or trained immunity via pattern-recognition receptors^21^. Some vaccines, such as the bacille Calmette-Guérin (BCG) tuberculosis vaccine, are hypothesized to initiate trained immunity^22^, and there is interest in further pursuing these targets in the field of tuberculosis and other infectious diseases. However, further study of the mechanisms behind this premature apoptosis in response to MAC biofilm is needed to identify specific molecular targets.

In Rose and Bermudez (2014), a reduction in apoptosis was accomplished *in vitro* by adding anti-TNF-R1 and anti-TNF-α^11^. However, in those experiments, the reduction in apoptosis also led to a significant increase in colony forming units (CFU), contrasting our result of reduction in bacterial load. We attribute these differences to different conditions of macrophage-bacterial interactions. In the lung airway, under our parameter ranges, we see a much higher macrophage to bacteria ratio and significantly less total biofilm mass, putting patrolling macrophages under significantly less apoptotic stress. Additionally, in our simulation and *in vivo*, infected macrophages can recruit additional macrophages from the vasculature, replenishing their numbers. Finally, Rose and Bermudez show high levels of apoptosis even with the anti-TNF-R1 or anti-TNF-α, indicating other pathways responsible, while we knocked out all apoptosis in response to biofilms. *In vitro*, it has been shown that TNF is associated with macrophages’ ability to kill MAC^23^. In our model, we mechanistically isolate the apoptotic response to biofilms from macrophages’ ability to kill bacteria, which is important, especially given the high proportions of intracellular bacteria that we observe at the end of our simulations. The mechanisms differentiating these two pathways need to be better understood to target apoptosis as a potential treatment, without compromising macrophages’ ability to kill MAC.

### 4.2 Reducing either frequency of bacterial exposure or bacterial load a patient is exposed to may have benefits

We show here that both bacterial load and bacterial deposition frequency can affect bacterial population counts and macrophage recruitment. Reducing either inoculum size or exposure frequency through environmental or patient lifestyle interventions may reduce likelihood of infection in at-risk patients. However, there does seem to be a threshold of frequency, around 2x/week, at which further reduction does not further decrease bacterial loads. Patient recommendations to reduce environmental exposure to MAC have included draining and refilling the household water heater every two weeks to reduce bacterial growth, cleaning showerheads frequently, and using filters on showers and tap water sources^24^, but none to this point have quantified frequency of exposure. Though we do not make a clinical recommendation based on this number, this indicates that further study or patient-centered exposure management or modifications may be of interest to clinicians.

### 4.3 Model Limitations

As in all models, we make simplifying assumptions due to parameter uncertainty and to limit model complexity. More detail can be added as biological mechanisms are identified and quantified, especially dose-response effects in macrophage-biofilm interactions. In fact, recent data suggest that patients with MAC-PD can have a dysregulated adaptive and possibly innate immunity, with exaggerated anti-inflammatory signals such as IL-10, to bacterial exposure^25^. Our current model does not distinguish between macrophage apoptosis and necrosis. Distinguishing between different types of cell death resulting from biofilm, bacterial or cytotoxic cell exposure would add further complexity to the system. Future studies on patient airway dynamics, such as quantitative cell counts of macrophages and chemokines profiles in bronchoalveolar lavage will serve as further validation data.

Further, our model focuses on dynamics within the lung airway, with limited movement in the z-dimension, and spatially dispersed bacteria. These dynamics may differ in a more confined environment such as lung lesions and nodules. Finally, we examine only the first two weeks of dynamics, and therefore assume no involvement of the adaptive immune system. Future model iterations over longer time periods will include the adaptive immune response and dynamics of nodules deeper in the lungs.

## 5. Study Highlights

### What is the current knowledge on the topic?

Though the incidence and prevalence of NTM infections, including MAC is growing, little is known about the role of their biofilms in overall infection. *In vitro* studies have shown that biofilms aid bacteria in providing protection from macrophage phagocytosis and antibiotics, while also causing premature macrophage apoptosis, but these dynamics have not been studied *in vivo*.

### What question did this study address?

What is the impact of MAC biofilms, host responses such as signaling, chemotaxis, apoptosis, and phagocytosis, and repeated bacterial exposure on tissue-level metrics such as bacterial load and phenotype distribution?

### What does this study add to our knowledge?

Biofilm indirectly allows for an increase in extracellular bacterial populations in the lungs by causing macrophage apoptosis, which in turn attracts more macrophages rather than allowing them to patrol the lung airway.

### How might this change drug discovery, development, and/or therapeutics?

This study explores the effects of specific targets in MAC infection while demonstrating the importance of spatiotemporal considerations in targeting host immune cells.

## 6. Acknowledgements

We thank Lev Gorenstein, Tsai-Wei Wu, and the rest of the Research Computing Staff for their assistance with batch computing at the Rosen Center for Advanced Computing. We thank Paige Marty, Renuka Reddy, Maleeha Shah, and Nianqiao Ju for their feedback throughout the research process. We also thank Alexa Petrucciani for feedback on figures and editing.

## 7. Author Contributions

C.W. wrote manuscript, designed research, performed research, analyzed data. K.P. designed research, provided clinical insights. P.E. designed research, provided clinical insights, provided funding. E.P. wrote manuscript, designed research, provided funding.

## 9. Supplement 1 – Parameter Table

**Table 1.** Parameters were taken from literature when possible. Parameter sources marked “e” indicates that the parameter was estimated, and given a large range due to uncertainty. Finally, diffusion coefficients were independently varied, but were calculated based on the diffusion rate of interleukin 6 (IL-6) in water^30^ divided by a value between 38 and 270 to account for the increased viscosity of mucus in diseases such as COPD and Cystic Fibrosis.

| Parameter Name | Description | Range | Unit | Source |
| --- | --- | --- | --- | --- |
| N_itlBac | Initial number of bacteria deposited | 1 – 50 | Scalar | e |
| F_itlPlank | Fraction of initial bacteria that are planktonic | 0.2 - 0.8 | Fraction | e |
| A_itlBiofilm | Amount of biofilm per initial sessile bacterium | 1 – 100 | Scalar | e |
| N_itlMac | Initial number of macrophages | 60 – 130 | Scalar | <sup>26</sup> |
| N_recAreas | Number of potential recruitment areas | 450-909 | Scalar | <sup>27</sup> |
| T_apopDeg | Time for dead macrophages to be removed/degrade | 0.5 – 72 | Hours | e |
| C_bacDeg | Fraction of total bacterial chemoattractant that degrades each tick | 0.0001 – 0.1 | %/timestep (6 min) | e |
| D_bacC | Bacterial chemoattractant diffusion coefficient | 1E-9 – 7E-9 | cm<sup>2</sup>/s | e |
| P_bacSurvApop | Probability of an individual IC bacterium surviving in apoptosis | 20 – 70 | % | e |
| P_bacInvPheno | Probability of an individual bacterium having an invasive phenotype | 6 – 15 | % | <sup>10</sup> |
| K_bacPerGridsquare | Bacterial carrying capacity per gridsquare | 20 – 300 | Scalar | e |
| T_ICBacDouble | Doubling time for IC bacteria | 100 – 140 | Hours | <sup>28</sup> |
| A_bacToC | Individual bacterium's contribution to bacterial chemoattractant per tick | 5 – 30 | scalar | e |
| M_bacZOffset | Maximum distance a child bacterium is offset from parent bacteria in z-dimension in extracellular replication | 0.001 | cm | e |
| M_bacXYRadius | Maximum distance child bacteria is offset from parent bacteria in xy-dimension in extracellular replication | 0.0006 – 0.0016 | cm | e |
| T_plankBacDouble | Doubling time for planktonic bacteria | 20 – 39 | Hours | <sup>29</sup> |
| P_phenoSwitch | Probability of an extracellular bacterium switching phenotypes given a bacterium of the opposite phenotype is in the same gridsquare, per tick | 0.01 – 0.1 | % | e |
| T_sessBacDouble | Doubling time for sessile bacteria | 40 – 80 | Hours | 29 |
| P_sessInv | Probability of a sessile bacterium in biofilm invading the epithelium | 0.001 – 0.1 | % | 10 |
| S_bacDiv | Variance in bacterial division | 10 | % | e |
| S_bacGrowth | Variance in bacterial growth rates | 10 | % | e |
| A_bacToBiofilm | individual sessile bacterium's contribution to biofilm per tick | 1E-6 – 1E-3 | Scalar | 11 |
| A_chemotaxThresh | threshold concentration of chemoattractant to bias macrophage phagocytosis | 1 | Scalar | e |
| P_infMPhag | probability of an infected macrophage phagocytosing a bacterium, per tick | 10 – 50 | % | e |
| N_threshBurst | number of IC bacteria necessary to cause an infected macrophage to burst | 10 – 60 | Scalar | 28 |
| C_macDeg | Fraction of total macrophage chemoattractant that degrades each tick | 0.0001 – 0.1 | %/6 min | e |
| D_macC | macrophage chemoattractant diffusion coefficient | 1E-9 – 7E-9 | cm<sup>2</sup>/s | e |
| A_infMToC | individual infected macrophage's contribution to macrophage chemoattractant per tick | 30 – 100 | Scalar | e |
| P_macKillsICBac | probability of infected macrophage killing an intracellular bacterium, per tick | 0.01 – 0.1 | % | e |
| P_mPhag | probability of healthy macrophage phagocytosing a bacterium, per tick | 50 – 100 | % | e |
| P_maxRec | maximum probability for recruitment of macrophages ($\beta$ of Hill function for recruitment) | 0.01-1 | % | e |
| P_halfMaxRec | half-max macrophage chemoattractant for recruitment of macrophages (K of Hill function for recruitment) | 20-1000 | Scalar | e |
| C_Nrec | Hill coefficient for variable recruitment rate | 2 | Scalar | e |
| areaDim | number of gridsquares in the x or y direction | 100 | Scalar |  |
| zDim | number of gridsquares in the z direction | 3 | Scalar |  |

## 10. Supplement 2 – Hill Function for Recruitment

**Supplement Figure 1.**
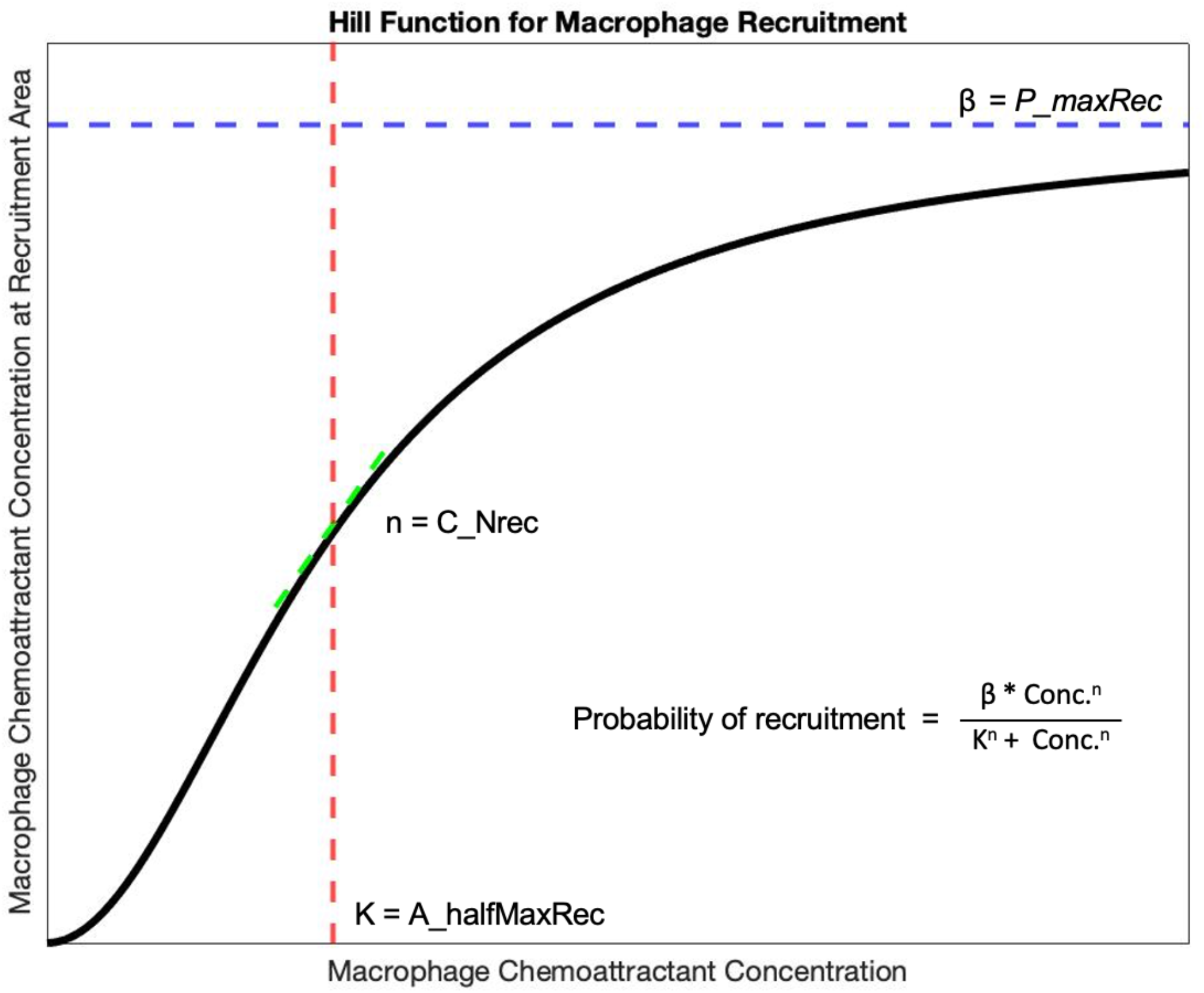
Graphical explanation of hill function parameters for recruitment. Probability of macrophage recruitment at each timestep is a function of chemoattractant concentration at that recruitment area, given by a hill function curve.

## 11. Supplement 3

**Supplement Figure 2.**
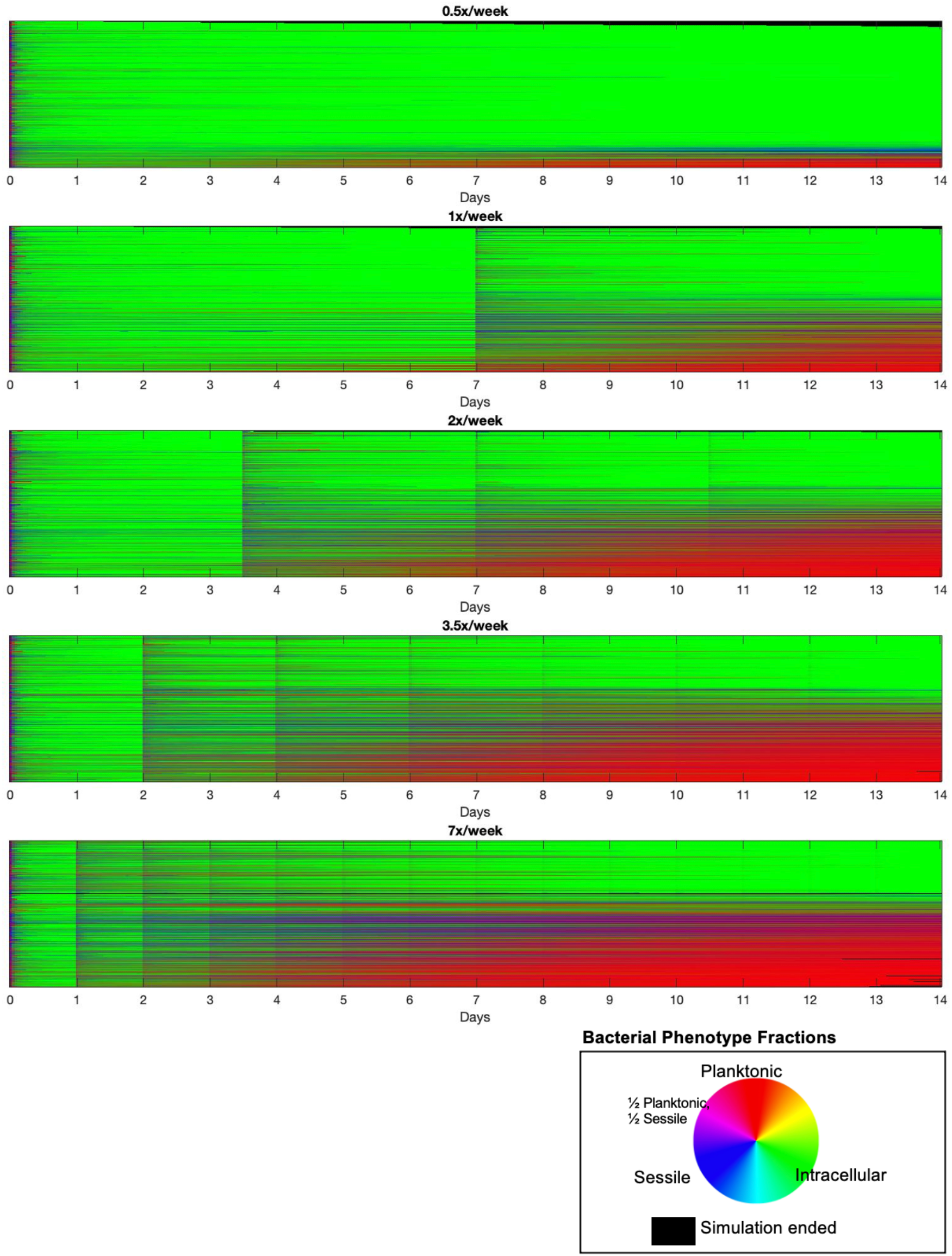
Bacterial phenotype distributions across deposition frequencies. We examine the relative bacterial proportions across all 900 simulations for each deposition frequency. The relative proportion of the bacterial phenotypes are plotted in color combinations of red, blue, and green, representing planktonic, sessile, and intracellular bacteria, respectively, as shown in the color wheel. Each horizontal line shows the distribution for one simulation. A black line shows simulations that ended after all bacteria were killed.

## 12. Supplement 4

**Supplement Figure 3.**
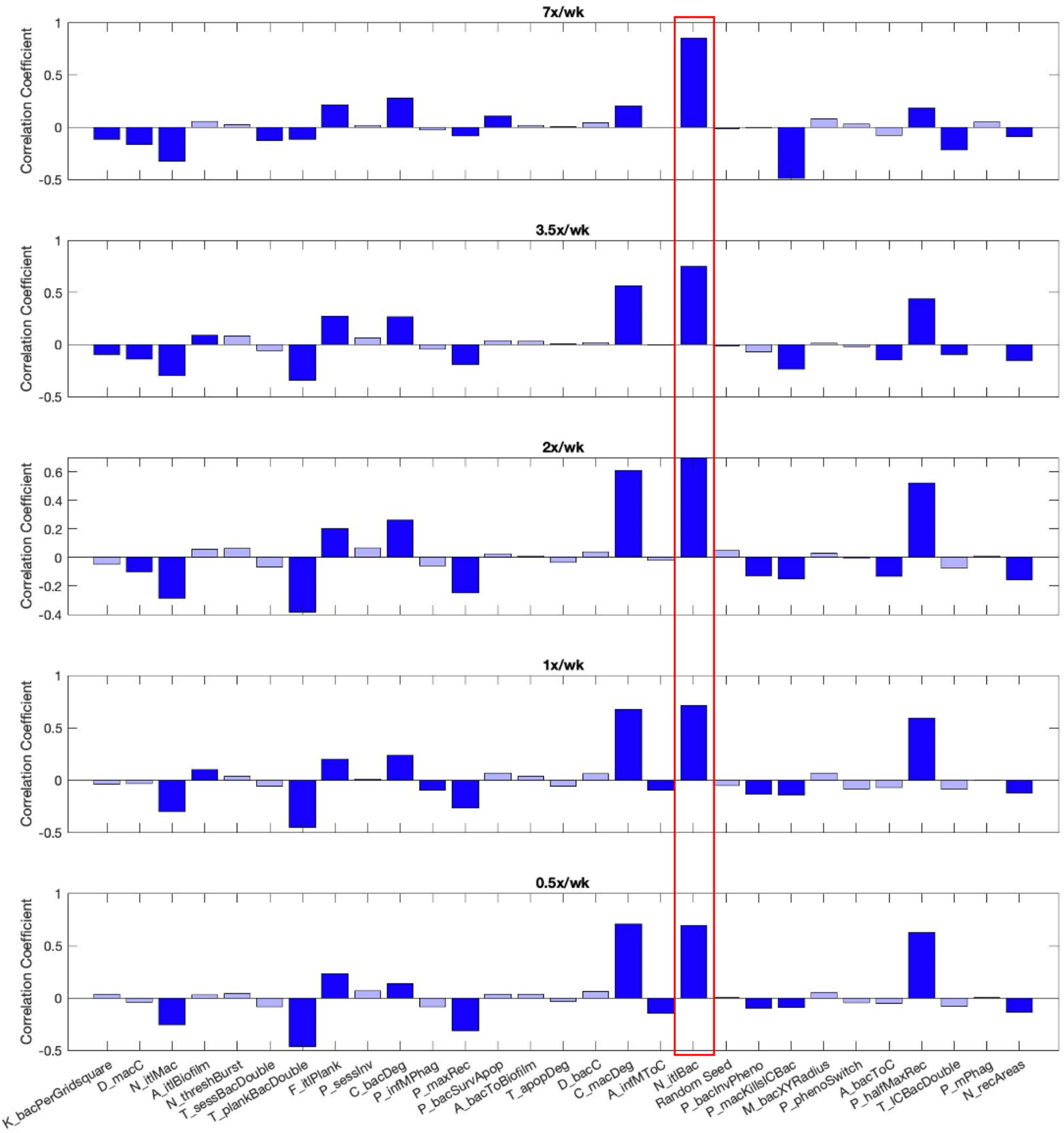
Partial Rank Correlation Coefficient (PRCC) for each parameter vs. final bacterial load across deposition frequencies. PRCC was calculated independently for all varied parameters in each deposition frequency. Significant correlation coefficients are shown in dark blue, while insignificant coefficients are in light blue. A red box is shown around *N_itlBac*, as it is consistently a driving parameter in bacterial load.

